# FapR regulates HssRS-mediated heme homeostasis in *Bacillus anthracis*

**DOI:** 10.1101/2024.07.08.602573

**Authors:** Hualiang Pi, Sophia M. Carlin, William N. Beavers, Gideon H. Hillebrand, Evan S. Krystofiak, Devin L. Stauff, Eric P. Skaar

**Affiliations:** Vanderbilt Institute for Infection, Immunology, and Inflammation, Vanderbilt University, Nashville, TN; Department of Pathology, Microbiology, & Immunology, Vanderbilt University Medical Center, Nashville, TN; Current address: Department of Microbial Pathogenesis and Microbial Sciences Institute, Yale University School of Medicine, New Haven, CT; Department of Biology, Grove City College, Grove City, PA; Department of Cell and Developmental Biology, Vanderbilt University, Nashville, TN

## Abstract

*Bacillus anthracis*, a Gram-positive facultative anaerobe and the causative agent of anthrax, multiplies to extraordinarily high numbers in vertebrate blood, resulting in considerable heme exposure. Heme is an essential nutrient and the preferred iron source for bacteria during vertebrate colonization, but its high redox potential makes it toxic in excess. To regulate heme homeostasis, many Gram-positive bacteria, including *B. anthracis*, rely on the two-component signaling system HssRS. HssRS comprises the heme sensing histidine kinase HssS, which modulates the activity of the HssR transcription factor to enable bacteria to circumvent heme toxicity. However, the regulation of the HssRS system remains unclear. Here we identify FapR, the transcriptional regulator of fatty acid biosynthesis, as a key factor in HssRS function. FapR plays an important role in maintaining membrane integrity and the localization of the histidine kinase HssS. Specifically, disruption of *fapR* leads to increased membrane rigidity, which hinders the penetration of HssRS inducers, resulting in the inactivation of HssRS. Furthermore, deletion of *fapR* affects the loading of HssS onto the cell membrane, compromising its heme sensing function and subsequently reducing endogenous heme biosynthesis. These findings shed light on the molecular mechanisms governing bacterial adaptation to heme stress and provide potential targets for antimicrobial intervention strategies.

**IMPORTANCE:** Understanding the mechanisms by which *B. anthracis* regulates heme homeostasis is crucial for developing new strategies to combat anthrax, a serious disease affecting both humans and animals. This study uncovers the role of the transcriptional regulator FapR in maintaining membrane integrity and facilitating the proper function of the HssRS two-component signaling system, which is essential for managing heme toxicity in *B. anthracis*, as well as other Gram-positive pathogens. By elucidating the connection between FapR and HssRS, our findings provide new insights into the molecular adaptation of bacteria to heme stress and expand our knowledge of bacterial physiology and pathogenicity. More importantly, targeting the regulatory pathways involved in heme sensing and homeostasis presents a promising approach for developing novel therapeutics against anthrax and potentially other bacterial infections that rely on similar mechanisms.

## INTRODUCTION

*Bacillus anthracis*, a zoonotic pathogen and Gram-positive, spore-forming bacillus, is notorious for its potential use as a bioterrorism weapon (1-4). It also causes endemic diseases, including cutaneous and gastrointestinal infections, in many regions worldwide (5). While *B. anthracis* spores are the infectious agent, the vegetative bacilli can cause extensive tissue damage through production of potent toxins and other virulence factors (3, 6, 7). Upon infection, spores are phagocytosed by innate immune cells and germinate into bacilli. The bacilli use a toxin-mediated mechanism to escape the host cell and proliferate to extreme densities in the bloodstream, leading to intoxication and death of the host (4).

As an intracellular pathogen, *B*. *anthracis* serves as an excellent model organism for studying stress responses in host environments. One important bacterial stress response mechanism is two-component systems (TCSs), which are crucial for detecting and adapting to various stimuli, including pH, temperature, nutrient scarcity, envelope integrity, and osmotic shock (8-14). The outcomes of TCS activation are diverse, influencing virulence, germination, sporulation, and antibiotic resistance (15, 16). TCSs are prevalent in bacteria but absent in humans and animals, rendering them promising targets for the development of antimicrobial therapeutics (17-19). On average, a bacterial genome contains about 30 TCSs. *B. anthracis* encodes 45 TCSs, highlighting the complex environmental challenges this pathogen encounters.

The heme sensing signaling system, HssRS, is a notable member of TCSs in Gram-positive bacteria (20-23). Heme, a vital nutrient and preferred iron source for certain bacteria, is contained within hemoproteins and accounts for 75% of the iron in vertebrates (24-28). Pathogens like *B. anthracis* have evolved to utilize heme from hemoglobin (24-27). This is particularly advantageous for *B. anthracis*, which can reach cell densities as high as 10^9^ colony forming units per milliliter of blood during systemic anthrax infection (4). However, heme also poses a toxicity risk by generating reactive oxygen species and inducing membrane stress (29). To manage heme homeostasis and mitigate heme-induced damage while still synthesizing, acquiring, and utilizing heme during infection (30), *B. anthracis* employs the HssRS signal transduction system to detect and regulate intracellular heme levels. Upon heme detection, the histidine kinase HssS undergoes autophosphorylation, subsequently activating its cognate response regulator HssR through phosphate transfer. Once phosphorylated, HssR initiates the expression of the heme-regulated ABC-type transporter HrtAB that effluxes excess heme from the cell. This transporter helps alleviate heme toxicity and promotes adaptation to heme exposure (22, 31-33).

In addition to heme, small molecules capable of triggering HssRS signaling have been identified, including the compound VU0038882 (‘882) (34). Exposure to ‘882 leads to an increase in endogenous heme biosynthesis, thereby activating HssRS signaling (34). Further investigation revealed that ‘882 specifically activates the coproporphyrinogen (CPG) oxidase, an enzyme that converts CPGIII to coproporphyrin III (CPIII) in the heme biosynthesis pathway (35), resulting in increased intracellular heme production. Notably, ‘882 exhibits toxicity toward bacterial cells undergoing fermentation by disrupting iron-sulfur cluster assembly (36).

The molecular mechanisms underlying the regulation of HssRS signaling to maintain heme equilibrium remain obscure. In this study, we employed a genetic selection approach to identify factors involved in modulating the HssRS system. We isolated spontaneous suppressor mutants, all with frame shift mutations in the *fapR* gene, which encodes the transcriptional repressor of fatty acid biosynthesis in Gram positive bacteria (37). These data demonstrate that disruption of *fapR* leads to complete inactivation of HssRS signaling triggered by ‘882 and partial inactivation of HssRS induced by heme. Utilizing a range of genetic, biochemical, and imaging techniques, these results elucidate important connections between membrane integrity, heme synthesis, and HssRS-mediated heme detection within the cellular membrane. These findings provide a foundation for developing antimicrobial strategies against this important pathogen.

## RESULTS

### Identification of regulatory factors involved in HssRS signaling

To identify regulatory factors required for HssRS activation, a genetic selection strategy was employed. A *B. anthracis* WT Sterne strain was engineered to contain two copies of *Escherichia coli relE*, driven by a *hssRS* promoter P*_hrt_* and integrated into two pseudogene loci (*bas3009*::P*_hrt_-relE bas4599*::P*_hrt_-relE*). This strain is designated as the 2xRelE strain (Fig. 1). The *relE* gene encodes an mRNA endoribonuclease that, upon activation of HssRS by ‘882 or heme, cleaves mRNA resulting in cell death (38). Colonies obtained from this selection represent strains with mutations impairing their ability to activate HssRS-dependent signal sensing and gene activation (Fig. 1A). The employment of two copies of *relE* prevents the selection of mutations within *relE* and enables preferential isolation of mutations that disrupt signal transduction of this TCS. Six suppressors were isolated from this RelE selection, all of which showed resistance to ‘882- or heme-mediated RelE killing and exhibited no mutations within *hssRS* (Fig. 1). Whole genome sequencing revealed that each isolated suppressor contained a frameshift mutation at one of three distinct locations within the same gene, *fapR* (Fig. 1 and Supplementary Table 3). No additional mutations were identified in these strains. These data indicate that FapR modulates HssRS activation in response to heme stress.

**Figure 1.**
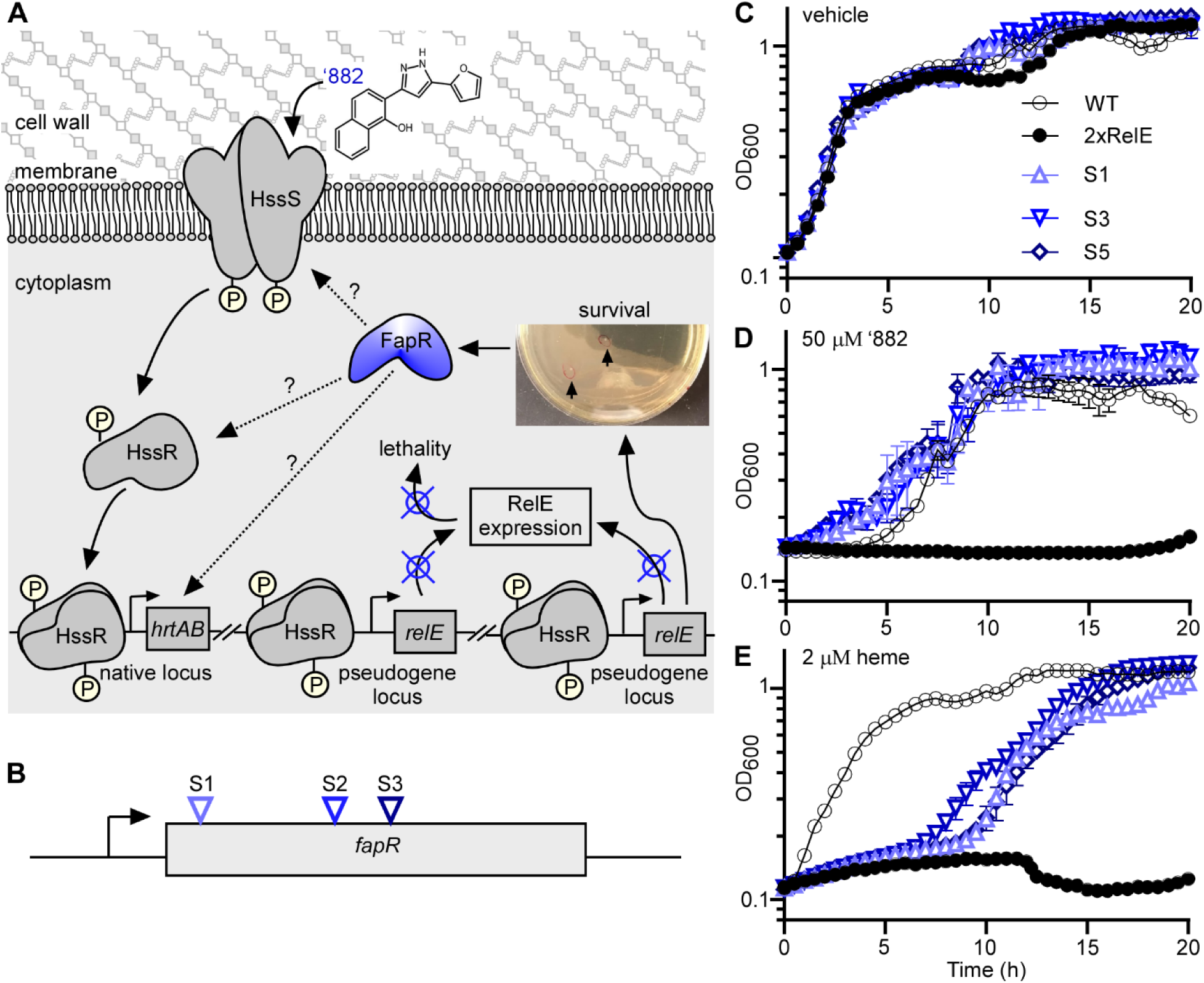
An unbiased genetic selection strategy identifies FapR as a regulatory factor in HssRS signaling. (**A**) Schematic of a genetic selection strategy: a 2xRelE strain (*bas3009*::P*_hrt_- relE bas4599*::P*_hrt_-relE*) was plated on medium containing activating compound ‘882 and colonies that arose represented bacteria that acquired mutations to inactivate the P*_hrt_* promoter. (**B**) All six suppressors isolated from genetic selections exhibit frame shift mutations at three different positions of *fapR*. Growth kinetics of *B. anthracis* WT, 2xRelE, and the representative suppressors in vehicle (**C**), 50 µM ‘882 (**D**), or 2 µM heme (**E**) was monitored for 24 h. Data shown are averages of three independent experiments (mean ± SD).

### Lipid membrane composition is altered by *fapR* deletion, which in turn suppresses ’882-mediated HssRS activation

FapR is a transcriptional repressor of fatty acid biosynthesis, largely conserved in Gram-positive bacteria (37). To validate the RelE genetic selection, we created a deletion mutant of *fapR* (*fapR*::*tet*). The loss of *fapR* results in overproduction of long-chain fatty acids (chain length of C_16_, C_17_, and C_18_) and a reduced production of shorter-chain fatty acids, particularly C_15_, in *Bacillus subtilis* (37). As anticipated, the *fapR* mutant showed a minor growth defect in rich medium (Fig. 2A), likely due to altered phospholipid synthesis. Interestingly, this mutant is highly resistant against ‘882 intoxication compared to WT. This phenotype can be complemented by providing *fapR in trans* (Fig. 2B).

**Figure 2.**
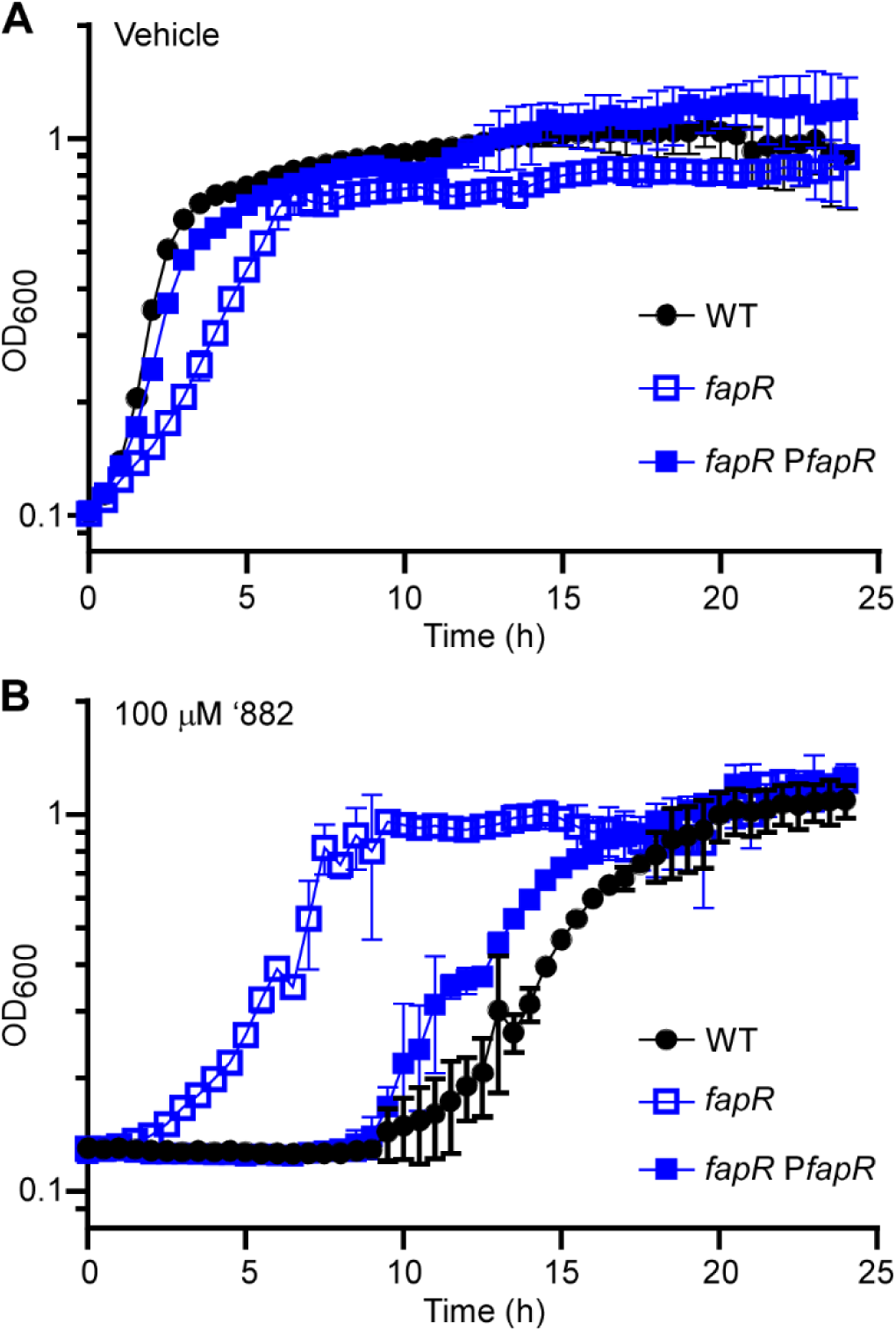
Deletion of *fapR* promotes tolerance against ‘882 intoxication in *B. anthracis*. The growth kinetics of *B. anthracis* Sterne WT, *fapR*::*tet* (*fapR*), and the complemented strain *fapR*::*tet* pOS1.P*_lgt_fapR* (*fapR* P*fapR*) grown in LB amended with (**A**) 0.1% DMSO (Vehicle) or (**B**) 100 µM ‘882. The *fapR* mutant is significantly more resistant against ‘882 toxicity. Data shown are averages of three biological replicates (mean ± SD) and experiments were repeated three times.

We hypothesized (i) the excessive synthesis of long-chain fatty acids caused by the deletion of *fapR* may lead to reduced membrane fluidity and increased membrane rigidity; and (ii) this alteration could hinder the penetration of ‘882 into the bacterial cell and diminish ‘882-mediated HssRS activation. To test this, we measured membrane fluidity using the fluorophore 1,6-diphenyl-1,3,5-hexatriene (DPH). DPH fluorescence reflects membrane fluidity or permeability as it only becomes fluorescent when embedded in membrane lipids. Indeed, the *fapR* mutant exhibited reduced membrane permeability in contrast to both the WT and *fapR* complemented strains (Fig. 3). This indicates that deletion of *fapR* leads to reduced membrane fluidity, consistent with previous findings in *B. subtilis* (37). This DPH assay was also repeated using the three representative suppressor mutants and similar results were observed (Fig. S1). Taken together, these results suggest that *B. anthracis* cells modify their lipid membrane composition through FapR, which prevents the infiltration of toxic compounds like ’882. Consequently, this modification, at least in part, contributes to the inability of ‘882 to stimulate HssRS signaling in the *fapR* mutant.

**Figure 3.**
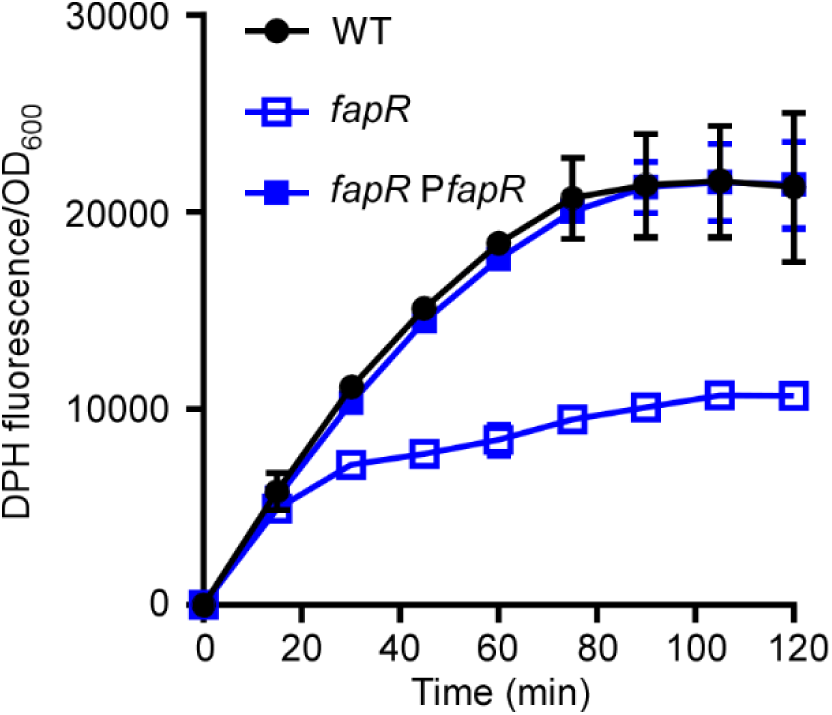
Deletion of *fapR* leads to decreased membrane permeability in *B. anthracis*. Membrane permeability was evaluated over 2 h in *B. anthracis* Sterne WT, *fapR*::*tet* (*fapR*), or the complemented strain *fapR*::*tet* pOS1.P*_lgt_fapR* (*fapR* P*fapR*) by monitoring the fluorescence of 1,6-diphenyl-1,3,5-hexatriene (DPH) (excitation, 365 nm; emission, 460 nm). All data represent means ± SD for measurements acquired in three biological replicates.

### Loss of *fapR* affects heme adaptation through HssRS

We then examined the impact of *fapR* deletion on heme mediated HssRS activation. Unlike the synthetic compound ‘882, which may passively enter cells without specific transporter systems, *B. anthracis* possesses a specific iron-regulated surface determinant (Isd) system for importing exogenous heme and an intrinsic heme synthesis pathway (30). This suggests that membrane permeability may affect HssRS activation triggered by heme differently. In *S. aureus*, heme biosynthesis is controlled post-transcriptionally by the enzyme responsible for the first step of this pathway, glutamyl-tRNA reductase (GtrR), which is regulated by both heme and the membrane protein HemX (39). Inactivation of *hemX* increases endogenous heme production which triggers HssRS-mediated upregulation of the heme exporter HrtAB, thereby effectively preadapting this mutant against heme toxicity in *S. aureus* (39). To test if this mechanism is conserved in *B. anthracis*, we created a *hemX* null mutant in *B. anthracis,* which exhibited no growth defects under vehicle treated conditions compared to WT (Fig. 4A). When grown in 20 µM heme, *B. anthracis* WT showed a noticeable growth defect with a 3.5-h lag phase (Fig. 4B). Preadaptation by pre-exposing cells to a low concentration of heme (2 µM) followed by exposure to toxic levels of heme (20 µM) enabled WT cells to develop robust resistance to heme toxicity (Fig. 4C). This preadaptation induces *hrtAB* upregulation through HssRS to alleviate heme intoxication, which was observed in all strains (Fig. 4C). In contrast, in the absence of preadaptation, the *hemX* mutant exhibited enhanced growth with no observable lag phase, while the *fapR* mutant showed elevated susceptibility to heme toxicity with prolonged lag phase (5 h) compared to WT (Fig. 4B-D). Intriguingly, the heme preadaptation phenotype observed in the *hemX* mutant is completely abolished in the *fapR hemX* double mutant (Fig. 4B-D). Together, these findings indicate that HemX regulates heme synthesis in *B. anthracis* similarly to *S. aureus*. Additionally, FapR specifically impacts HssRS activation stimulated by endogenous heme biosynthesis.

**Figure 4.**
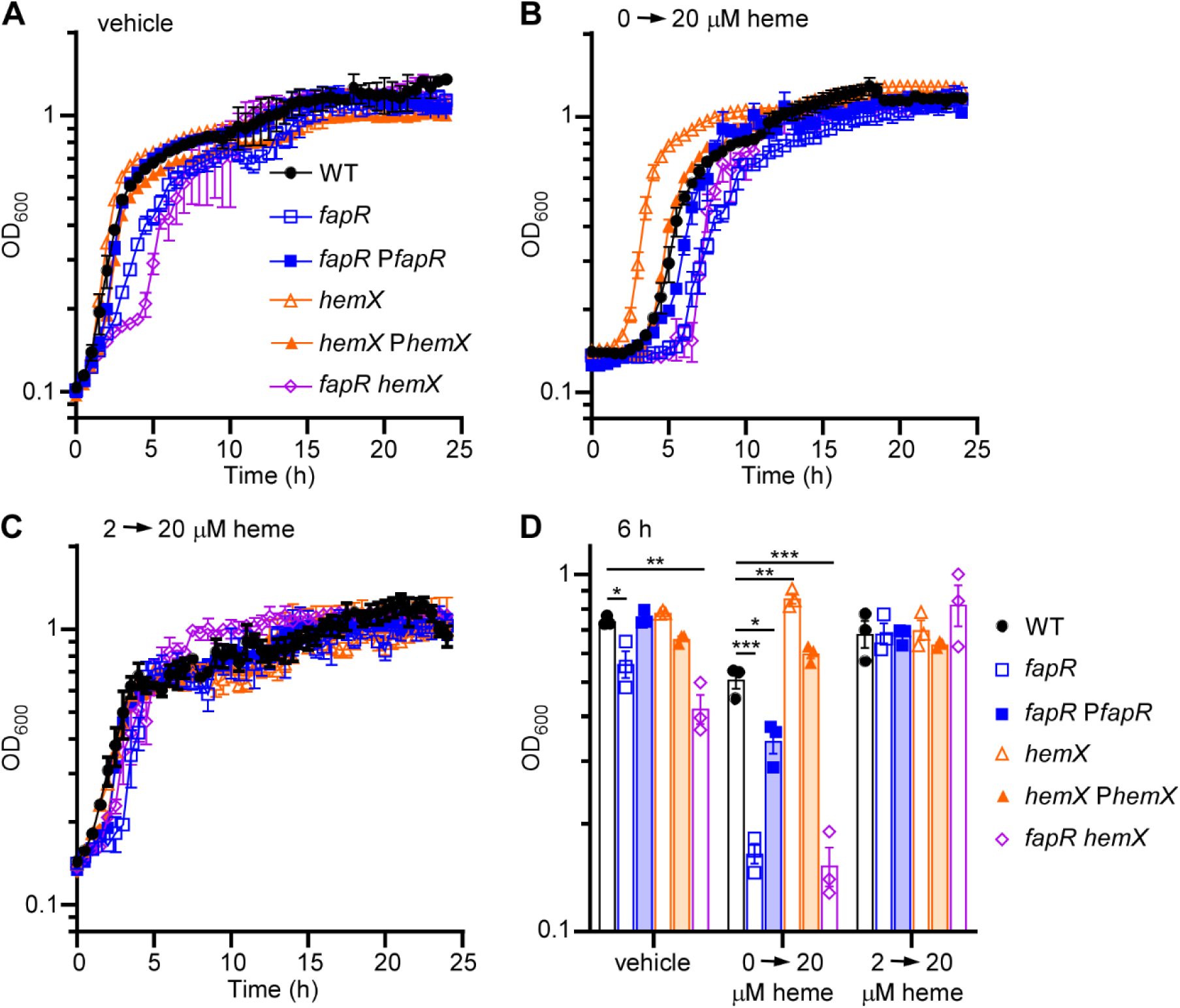
Deletion of *hemX* enables *B. anthracis* to pre-adapt to heme intoxication, a capability that is abolished in the *fapR hemX* double mutant. Growth kinetics of *B. anthracis* WT, *fapR*, Δ*hemX*, the complemented strains *fapR*::*tet* pOS1.P*_lgt_fapR* (*fapR* P*fapR*) and Δ*hemX* pOS1.P*_lgt_hemX (hemX* P*hemX*), and *fapR hemX* in (**A**) vehicle or (**B**-**C**) 20 µM heme was monitored for 24 h. Prior to the measured growth, the strains were pre-grown to the early stationary phase in LB medium containing (**B**) vehicle or (**C**) 2 µM heme. (**D**) Growth as measured by OD_600_ at 6 h under varied conditions. All data represent means ± SD for measurements acquired in three biological replicates.

### The deletion of *fapR* disrupts both HssS localization and cell morphology

Deletion of *fapR* abolishes the resistance of *B. anthracis hemX* null mutant to heme toxicity (Fig. 4). We hypothesized that in the *fapR* mutant, increased membrane rigidity disrupts the localization of the histidine kinase HssS within the membrane, consequently leading to dysregulated heme homeostasis. To evaluate HssS localization, we created a protein fusion construct, HssS-GFP (pOS1.P_lgt_*hssS*.*gfp*). In WT cells, the GFP signal exhibits a uniform distribution across the membrane (Fig. 5A-C). However, in the *fapR* deletion mutant, distinct GFP puncta are observed at division planes, accompanied by noticeable GFP signal diffusing throughout the cytosol (Fig. 5D-F). Although the total GFP signal relative to the DNA signal remains comparable between the two strains, WT shows significantly lower GFP signal in the cytosol compared to the *fapR* mutant (Fig. 5G). Together, these data indicate that the deletion of *fapR* disrupts HssS loading onto the cell membrane.

**Figure 5.**
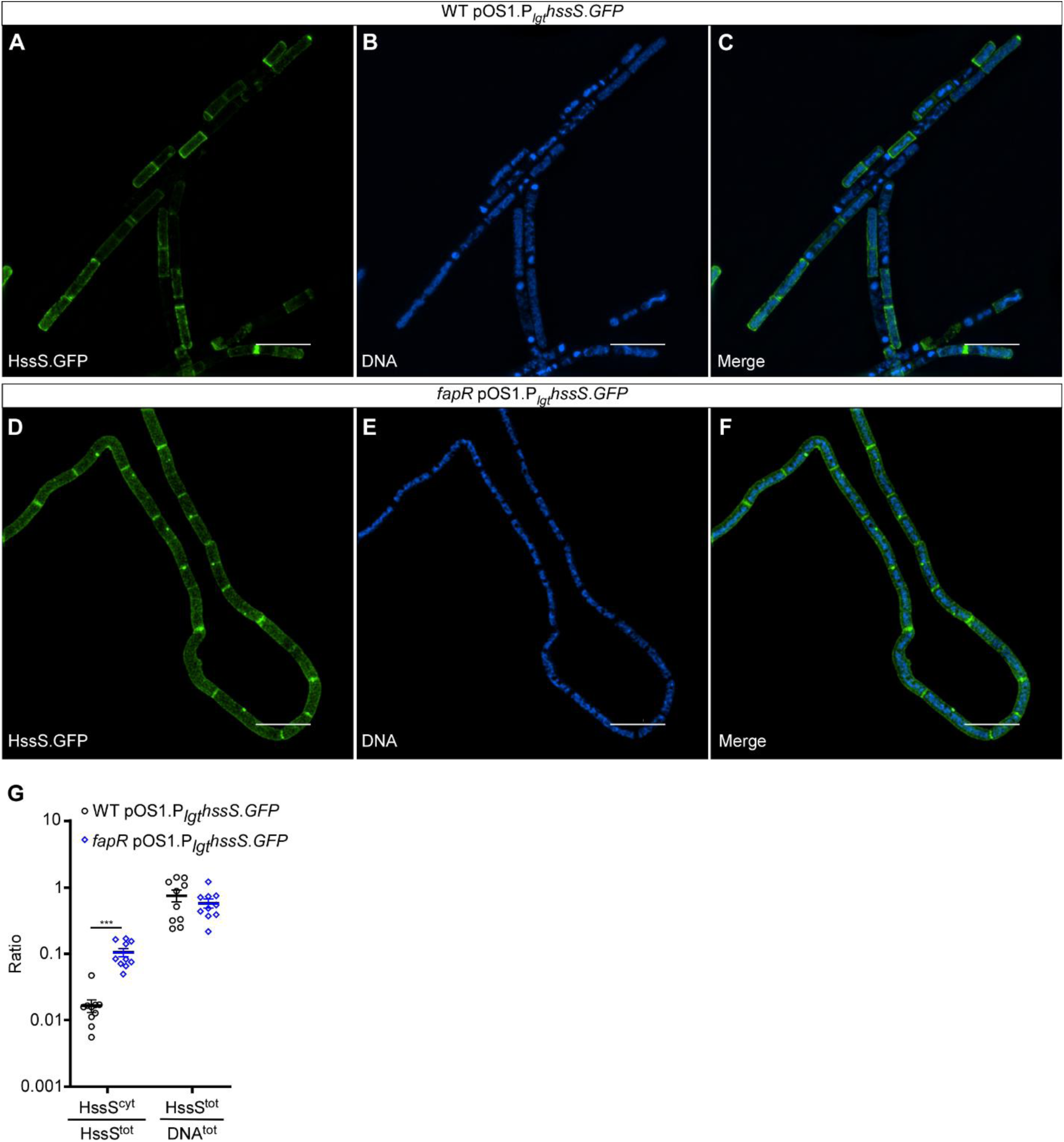
Deletion of *fapR* disrupts HssS membrane localization. (**A**-**F**) Representative super-resolution images depicting HssS localization in (**A**-**C**) WT or (**D**-**F**) *fapR* harboring pOS1.P_lgt_*hssS*.*GFP*. Cells were grown to OD_600_∼1 prior to staining with Hoechst. (**G**) HssS^cyt^/HssS^tot^, ratio of cytoplasmic HitS signal intensity (HssS signal overlapping with DNA signal) relative to total HssS signal intensity; HssS^tot^/DNA^tot^, total HssS signal intensity normalized to total DNA signal. Data shown are averages of 10 individual images collected per sample from two individual experiments (mean ± SEM). Statistical significance was determined using a two-way ANOVA (*, P ≤ 0.05; **, P ≤ 0.01; ***, P ≤ 0.001). Scale bar, 5 mm.

Furthermore, we noticed that WT cells typically form short, regular chains, whereas the *fapR* mutant cells exhibit a range of morphological abnormalities, including bending, coiling, and filamentation (Fig. 5 and Fig. S2). Although septa are visible in both strains, the *fapR* mutant cells frequently fail to separate. To further characterize this morphological defect, we examined the surface structures of both strains using scanning electron microscopy (SEM). The elongated filaments produced by the *fapR* mutant cells are intertwined to form bundles, with some displaying pronounced coarse ruffling patterns, unlike the smooth surface of WT cells (Fig. 6). This aberrant morphology suggests a potential correlation with the observed membrane rigidity in this mutant. This rigidity, in turn, interferes with HssS membrane localization and heme sensing through HssRS.

**Figure 6.**
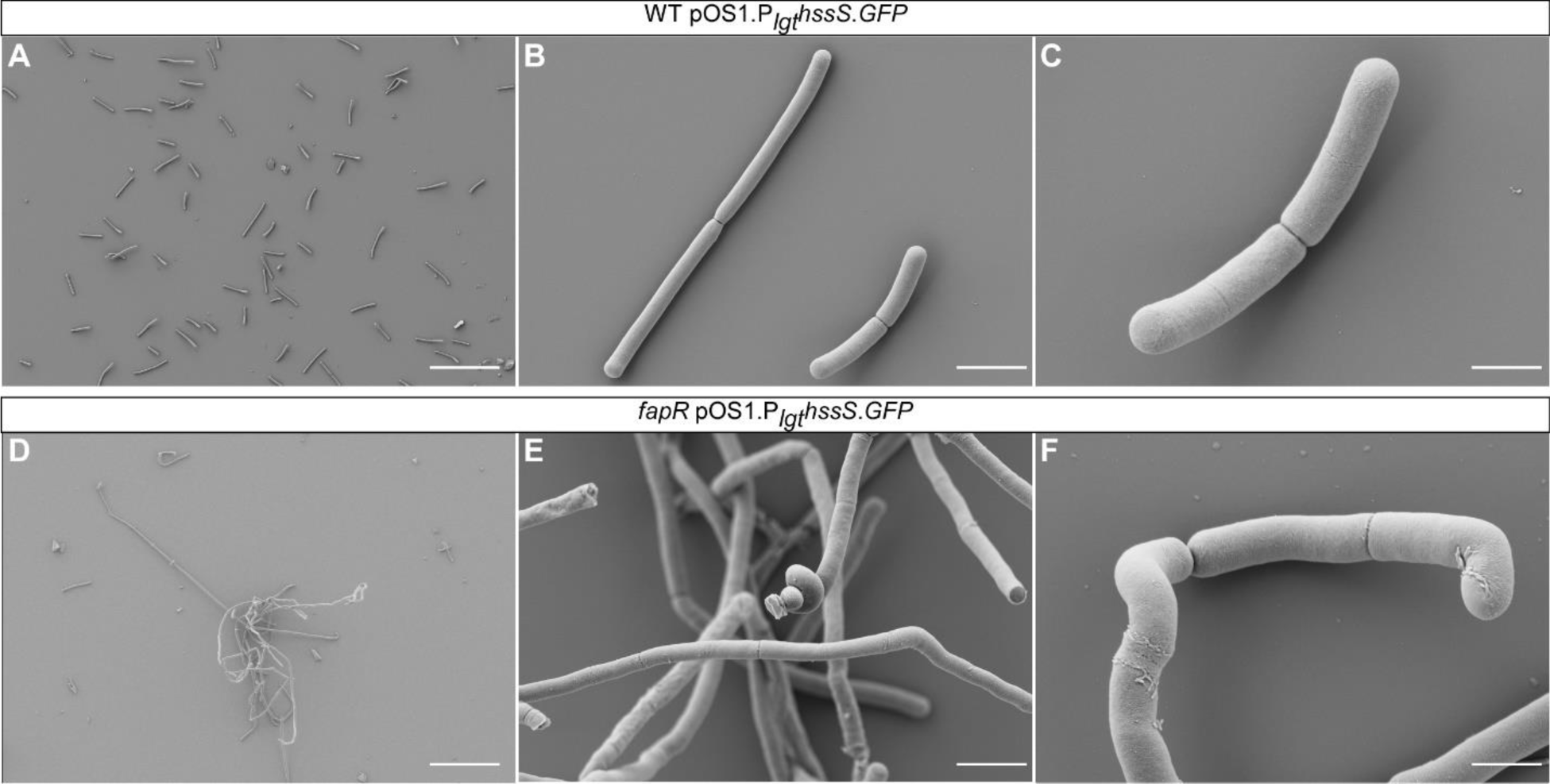
Deletion of *fapR* alters cell morphology of *B. anthracis*. Representative scanning electron microscopy (SEM) images of WT (**A**-**C**) and *fapR* (**D**-**E**) strains harboring pOS1.P_lgt_*hssS*.*GFP* are shown. Scale bars, 30 µm (**A**, **D**), 3 µm (**B**, **E**), and 1 µm (**C**, **F**).

### Deletion of *hemX* activates HssRS but this activation is compromised in the *fapR* mutant

Next, we evaluated the impact of *fapR* and *hemX* deletions on HssRS signaling using a XylE reporter. The expression of *xylE* is driven by the *hrt*AB promoter (P*_hrt_*), making XylE activity a proxy for the P*_hrt_* promoter activation regulated by the HssRS system. Under vehicle conditions, the basal level of promoter activity is significantly lower in the *fapR* mutant compared to WT. In contrast, the *hemX* mutant shows hyperactive promoter activity, which is significantly reduced by the deletion of *fapR* in the double mutant (Fig. 7A). This indicates that excess endogenous heme synthesis in the *hemX* mutant activates the heme stress response, similar to what has been observed in *S. aureus* (39). However, this activation is diminished by the deletion of *fapR* (Fig. 7A). Both ‘882 and the heme intermediate CPIII activate the HssRS system, although not as strongly as heme (Fig. 7B-D). Notably, *fapR* deletion affects the activation of the HssRS system under all tested conditions, including exogenous heme treatment (Fig. 7B-D).

**Figure 7.**
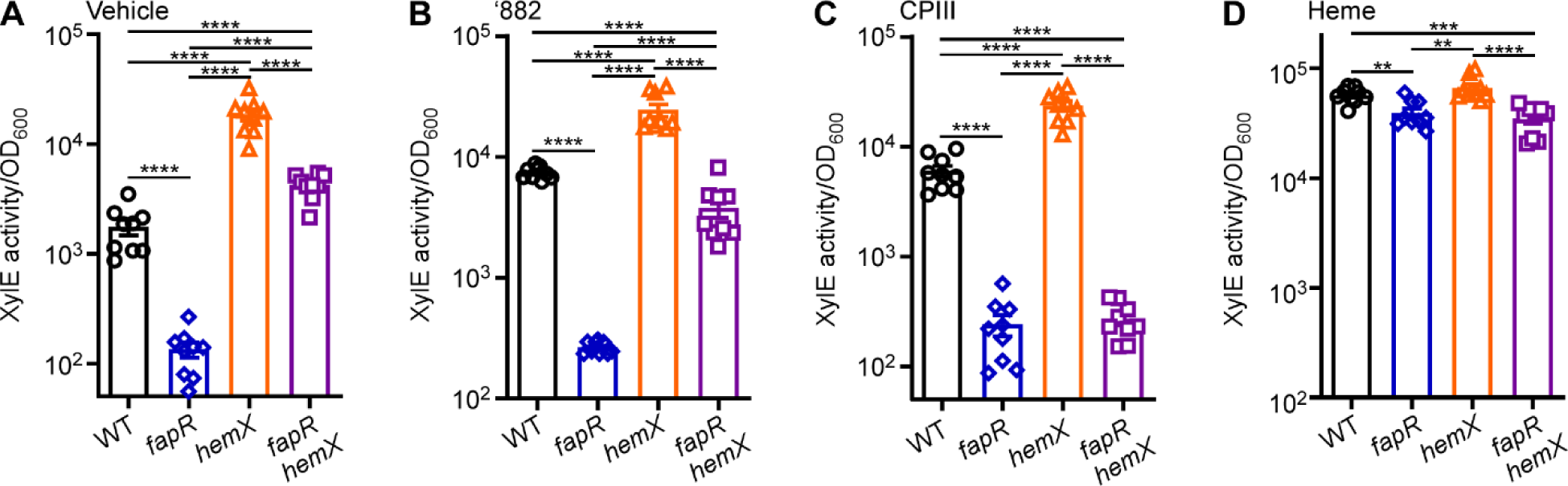
The heme sensing activity is hyperactive in the *hemX* mutant, which is compromised by *fapR* deletion. *B. anthracis* strains harboring pOS1. P*_hrt_*-*xylE* growing in LB to mid-logarithmic phase, and (**A**) vehicle, (**B**) 25 µM ‘882, (**C**) 10 µM CPIII, and (**D**) 10 µM heme was amended. After additional 14 h incubation at 37 °C, cell cultures were harvested and the XylE activity was quantified. All data are mean ± SEM (n=9). Statistical analyses were done using one-way ANOVA ∗P < 0.05, ∗∗P < 0.01, ∗∗∗P < 0.001, and ∗∗∗∗P < 0.0001.

To further validate these results, we employed an mCherry reporter (P*_hrt_*-*mCherry*). Given the hyperactive promoter activity in the *hemX* mutant, the fluorescence signal in this strain instantly reached its maximum beyond the detection limit, resulting in an “overflow signal”. Thus, the time taken to reach this overflow signal serves as a proxy for P*_hrt_* promoter activation regulated by the HssRS system (Fig. S3) with shorter times equating to higher promoter activity. Inactivation of *fapR* significantly affects the promoter activity of the HssRS system under all conditions tested, including heme treatment (Fig. S3). Together, these data suggest that the deletion of *fapR* impairs the ability of HssRS to sense endogenous heme, while its detection of exogenous heme is significantly affected but still largely intact.

### Inactivation of *fapR* reduces endogenous heme synthesis

The distinct response of the HssRS system to exogenous and endogenous heme is intriguing, particularly given that FapR is exclusively required for activating HssRS signaling through endogenous heme. This raises the possibility that FapR may also play a role in heme biosynthesis. To explore this, we quantified the intracellular levels of heme and its intermediate, CPIII using liquid chromatography-tandem mass spectrometry (LC-MS/MS). The *fapR* mutant exhibited diminished production of both heme and CPIII, while the *hemX* mutant showed significantly elevated levels of both compared to WT (Fig. 8), consistent with a prior study in *S. aureus* (39). Notably, the CPIII level in the double mutant fell between those of the WT and the *hemX* mutant, but the heme level in the double mutant was even lower than that in the WT (Fig. 8) for reasons that remain unclear. Taken together, these data indicate that FapR affects heme mediated HssRS activation through its regulatory control of membrane integrity and endogenous heme synthesis.

**Figure 8.**
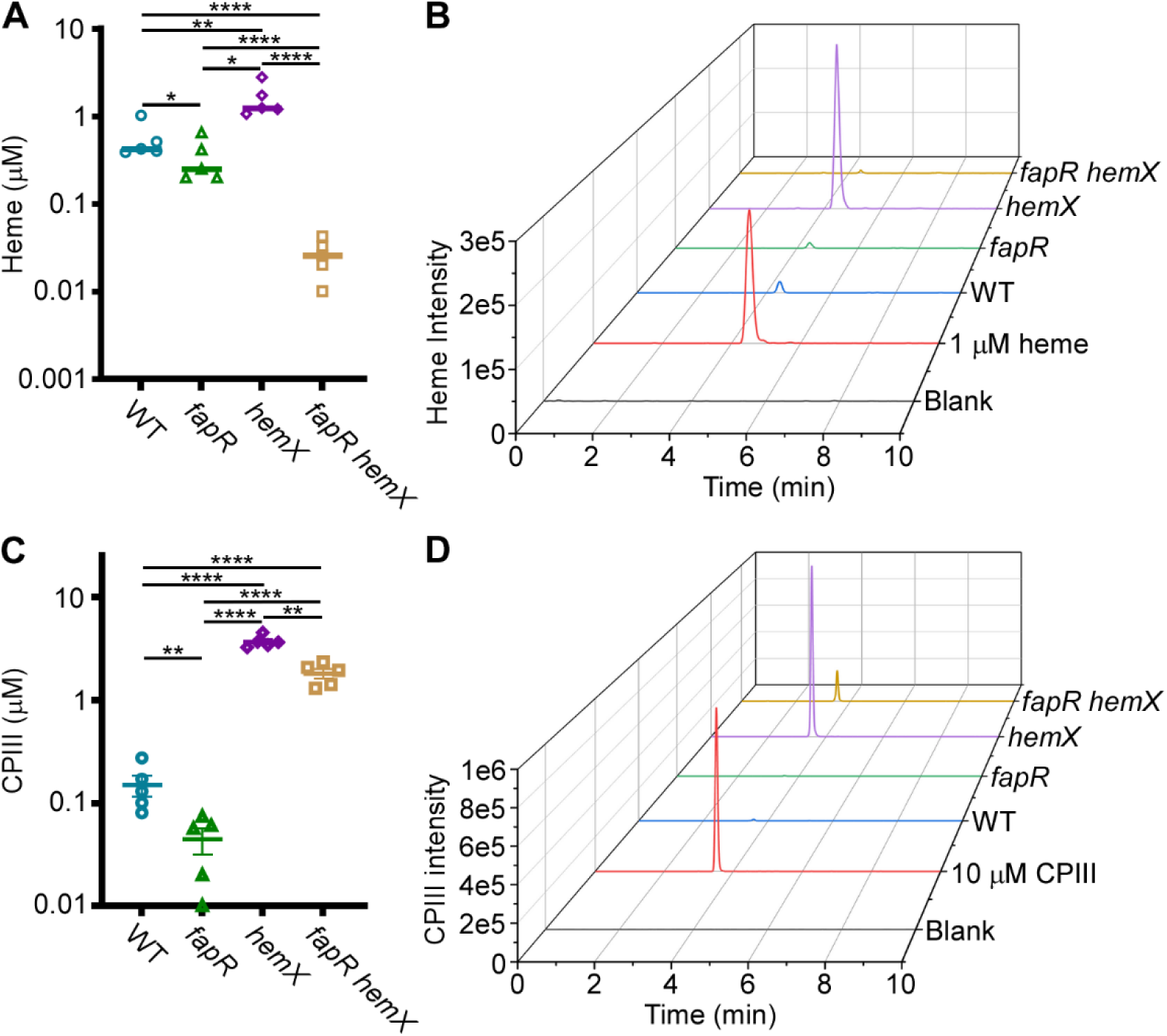
Inactivation of *fapR* reduces heme synthesis while deletion of *hemX* increases this process. (**A**, **B**) Heme and (**C**, **D**) coproporphyrin III (CPIII) were quantified in *B. anthracis* Sterne WT, *fapR*::*tet*, *hemX*, *fapR hemX*, and *hssRS*. All data are mean ± SD (n=5). Statistical analyses were done using one-way ANOVA ∗P < 0.05, ∗∗P < 0.01, ∗∗∗P < 0.001, and ∗∗∗∗P < 0.0001. Representative graphs of (**B**) heme and (**D**) CPIII are shown.

## DISCUSSION

Proper synthesis of the cell membrane and maintenance of membrane integrity are vital for bacterial fitness. Under stressful conditions and chemical insults, it becomes critical to modulate membrane fluidity to block toxic compound from entering and ensure cell survival. This study uses *B. anthracis* as a model organism to investigate the regulatory mechanism of the HssRS signaling system. The findings illustrate that bacterial cells employ an extreme tradeoff strategy: disrupting the FapR regulator of fatty acid synthesis leads to various fitness defects, including compromised membrane fluidity and abnormal cell morphology (Figs. 5-6 and S2), but these changes counteract harmful chemicals such as the HssRS activating compound ‘882.

The synthetic compound ‘882, structurally distinct from heme, does not activate the HssRS heme sensor through direct ligand binding (34). Instead, ‘882 activates the essential heme synthesis enzyme CgoX, leading to the accumulation of endogenous heme and its precursor CPIII. These compounds, in turn, trigger the HssRS signaling system, which activates the heme-regulated ABC-type transporter HrtAB to mitigate heme intoxication, as demonstrated previously (35) and in this study (Figs. 7 and S3). The chemical properties of ‘882 suggest it is membrane permeable (40, 41), and *B. anthracis* likely does not encode specific transporters for this synthetic compound. Therefore, the most effective way to prevent ‘882 intoxication is by blocking its entry into the cells through modulating the membrane fatty acid composition via FapR.

In addition to ‘882, the HssRS system detects heme, both endogenous and exogenous (22, 31, 32, 42) (Figs. 4, 7, and S3), though the specific molecular basis underlying HssS heme sensing remains largely obscure. Recent work, utilizing Alphafold2 prediction and structural simulations, has revealed a heme-binding site situated within the HssS dimer, with several conserved residues in this pocket essential for HssS activation (43). The heme binding pocket is located at the interface between the cell membrane and HssS extracellular domains (43). The authors propose a model wherein heme, either extracellular or intracellular, passively enters the membrane lipids to activate HssS at a common site within the cell membrane (43). However, the mechanisms by which HssS discriminates between endogenous and exogenous heme are unclear, as are the effects of lipid composition and membrane fluidity on HssS heme detection. Our data demonstrate that deletion of *fapR* leads to a significant reduction in HssS activation by exogenous heme, although the overall HssS activity remains relatively unaffected (Figs. 7 and S3). This could be due to increased membrane rigidity caused by *fapR* deletion, which compromises the proper loading of HssS onto the cell membrane (Figs. 3 and 5), resulting in fewer functional HssS proteins available to precisely manage intracellular heme homeostasis. Consequently, cells become less responsive under heme stress. Further structural and biochemical characterization is needed to elucidate the HssS detection mechanism of both endogenous and exogenous heme.

The HssRS signaling system is essential for heme adaptation (42, 44). In the presence of sub-toxic heme concentrations, bacterial cells upregulate the HrtAB efflux transporter through HssRS, enabling them to adapt and tolerate increased heme levels. This finely tuned activity of HssRS ensures that cells resolve the heme paradox by maintaining intracellular heme concentrations at necessary but non-toxic levels (22, 31, 32, 42). To examine the effects of *fapR* deletion on heme adaptation capacity, we employed a genetic tool using the *hemX* deletion mutant. In this mutant, elevated endogenous heme levels activate HssRS, enabling cells to effectively preadapt to heme toxicity, as demonstrated previously in *S. aureus* (39) and now in *B. anthracis* (Fig. 4). Notably, this preadaptation is completely abolished in the *fapR hemX* double mutant (Fig. 4). The explanation for this is twofold. First, as stated earlier, by reducing membrane fluidity, the deletion of *fapR* leads to a decrease in the number of functional HssS proteins capable of effectively regulating heme equilibrium (Figs. 3 and 5). Second, the deletion of *fapR* results in reduced levels of both heme and its precursor CPIII (Fig. 8), further repressing HssRS signaling activity and eliminating the preadaptation phenotype observed in the *hemX* mutant. However, the specific enzyme(s) in the heme biosynthesis pathway affected by FapR remain unidentified. Ongoing studies aim to uncover the molecular mechanism of this regulation. Collectively, this study highlights FapR as a modulator of the HssRS system, affecting HssRS activation through its regulatory control of membrane integrity and heme biosynthesis. Additionally, these results establish a link between fatty acid synthesis and heme detection through the HssRS signal transduction system.

## MATERIALS AND METHODS

### Bacterial strains and growth conditions

All strains and plasmids used in this study are listed in Table S1. Cells were grown in LB medium with shaking at 180 RPM or on solid LB agar plates with appropriate antibiotic selection at 37°C. The antibiotics concentrations used were: carbenicillin (carb, 50 µg ml^-1^), chloramphenicol (cam, 30 µg ml^-1^), kanamycin (kan, 40 µg ml^-1^), and erythromycin (erm, 20 µg ml^-1^). Stocks of the compound ‘882 (100 mM) were prepared in DMSO and stored at -20°C.

### DNA manipulation and strain construction

To generate a plasmid for deletion of *fapR* or *hemX*, flanking sequences were inserted into the mutagenesis plasmid pLM4(45). Briefly, flanking sequences were amplified using a distal primer containing a restriction enzyme site found in pLM4 (XmaI or SacI) and a proximal primer containing a short overlapping sequence. The PCR-amplified DNA was fused using PCR sequence overlap extension (PCR-SOE) as described previously (46) and inserted between the XmaI and SacI sites in pLM4, generating the plasmid pLM4-*fapR* or pLM4-*hemX*. Genetic manipulation in *B. anthracis* using pLM4-*fapR* or pLM4-*hemX* was performed as previously described (16) to produce *B. anthracis fapR*::*tet* or *hemX* null mutants. All strains used in this study were verified by PCR using primers listed in Table S2.

### Genetic selection

To isolate spontaneous suppressor mutants that inactivate HitRS signaling, we utilized a 2xRelE strain (*bas3009*::P*_hrt_*-*relE bas4599*::P*_hrt_*-*relE*) (38). The 2xRelE strain was streaked onto an LB agar plate and single colonies were grown in 5 ml LB amended with 50 µM ‘882 for 18 hours at 37°C with vigorous shaking. Serial dilutions of the bacterial cultures were prepared, and 100 μl of the 10^-5^ dilution was plated onto BHI agar plates containing 25 mM glucose and 100 µM ‘882. The plates were incubated at 37°C for up to 2 days. Spontaneously arising colonies were then isolated, streaked onto fresh LB agar plates for single colonies, and saved for further analysis.

### Whole genome sequencing

Genomic DNA (gDNA) was extracted from the suppressor mutants using the Qiagen DNeasy Blood and Tissue kit, following the manufacturer’s instructions. The purified gDNA was sequenced by Genewiz using the Illumina NextSeq 550 platform with 150 Mbs coverage. Sequencing reads were trimmed and mapped to the genome of *B. anthracis* Sterne (T00184), and variant detection was performed on the mapped reads using CLC Genomics Workbench v20.0.1 with default settings.

### Growth curves

Cells were grown overnight in LB medium at 30°C, subcultured at a 1:100 ratio into fresh LB medium and incubated for 6 h at 37°C with vigorous shaking. Cell density (OD_600_) was then monitored every 30 min for 24 h at 37°C with continuous shaking using a BioTek Epoch2 spectrophotometer. Experiments were conducted at least three times with three biological replicates per experiment. Data shown are averages of three replicates (mean ± SD).

### Membrane permeability assay

The membrane permeability of various *B. anthracis* strains was examined using 1,6-diphenyl-1,3,5-hexatriene (DPH), which fluoresces exclusively within the lipid membrane (47). Briefly, cells were grown overnight in LB medium at 30°C, subcultured at a 1:100 ratio into fresh LB medium, grown for 6 h at 37°C with vigorous shaking before being diluted tenfold. Cell density (OD_600_) and fluorescence (excitation, 365 nm; emission, 460 nm) were then monitored every 20 min for 2 h at 37°C with continuous shaking using a BioTek fluorescence plate reader. Each experiment was conducted at least three times with three biological replicates each time. The presented data are averages of the three replicates (mean ± SD).

### Structured illumination microscopy (SIM) imaging

*B. anthracis* strains containing the HssS-GFP protein fusion vector were grown overnight in LB medium. Cultures were harvested and washed in PBS. DNA staining was performed using 1 mg ml^-1^ Hoechst 33342 (ThermoFisher) for 10 min at room temperature. Subsequently, cells were fixed in 4% paraformaldehyde (Invitrogen) for 30 minutes at room temperature. Following fixation, the cells were washed with PBS once and then with water twice before being dried onto a coverslip. The prepared coverslips were mounted to slides using Prolong Gold (ThermoFisher). Super resolution imaging was conducted using Nikon SIM. Image collection settings were calibrated using WT control samples. Brightness and contrast were adjusted to visualize both DNA and membrane localization of the GFP-tagged HssS. Analysis was carried out with Nikon Elements General Analysis 3 (GA3) software. Signal thresholds were determined for each channel, and for each signal, the sum intensity, total volume, overlapping volume, and overlapping signal intensity were calculated using GA3.

### Scanning electron microscopy (SEM) imaging analysis

*B. anthracis* cells grown in LB medium were washed with PBS buffer and fixed overnight at 4°C in 0.05 M sodium cacodylate buffer containing 2.0% paraformaldehyde and 2% glutaraldehyde. Samples were postfixed in 1% OsO4 and further stained in 1% uranyl acetate. The samples were progressively dehydrated in a graded ethanol series, followed by critical point drying using a Tousimis Samdri PVT-3D. The dried samples were mounted on SEM stubs and sputter-coated with platinum using a Cressington 108. To facilitate grounding and prevent charging during SEM imaging the edges of the coverslips were painted with conductive carbon paint (Electron Microscopy Sciences). Scanning electron microscopy was performed using a Zeiss Crossbeam 550 FIB-SEM at 2 keV using the lower secondary electron detector.

### XylE reporter assay

To examine the effects of *fapR* or *hemX* deletion on HssRS activation, an XylE reporter construct (P*_hrt_*-*xylE*) was generated to evaluate the promoter activity. Cells were grown overnight at 30°C in LB medium and then subcultured at a 1:100 ratio into fresh LB medium. After 18 h of growth at 37°C, in the presence or absence of the specified HssRS activator, the abundance of the XylE enzyme (catechol 2, 3-dioxygenase) in *B. anthracis* cell lysates was quantified. This was done by measuring the rate at which catechol was converted to 2-hydroxymuconic acid using a spectrophotometer.

### mCherry reporter assay

To further validate the effects of *fapR* or *hemX* deletion on HssRS activation, an mCherry reporter construct (P*_hrt_*-*mcherry*) was generated to examine the promoter activity. *B. anthracis* strains harboring pOS1 P*_hrt_*-*mcherry* were grown overnight at 30°C in LB medium and subcultured at a 1:100 ratio into fresh LB medium. These cultures were then inoculated in LB amended with varied compound as indicated. Cell density (OD_600_) and fluorescence (Ex: 590 nm; Em: 620 nm) were then monitored every 30 min for 20 h at 37°C with continuous shaking using a BioTek fluorescence plate reader. The time taken to reach an overflow signal, indicating maximum fluorescence beyond the detection limit, serves as a proxy for P*_hrt_* promoter activation regulated by the HssRS system. Each experiment was conducted at least three times with three biological replicates each time. The presented data are averages of the three replicates (mean ± SD).

### Heme quantification

Samples for quantification of heme and heme biosynthetic intermediates were analyzed on a Thermo TSQ Quantum Ultra with an ESI source interfaced to a Waters Acquity UPLC system in the Vanderbilt Mass Spectrometry Research Center. Analytes were separated by gradient HPLC with an Agilent Poroshell 120 EC-C18 C_18_ (3 x 50 mm, 2.7 μm) and a Phenomenex SecurityGuard C_18_ cartridge (3.2 x 8 mm) at a flow rate of 0.4 mL/min using 0.1 % formic acid in water and 0.1 % formic acid in acetonitrile as the A and B mobile phases, respectively. The gradient was held at 25 % B for 0.5 min, then ramped to 100 % B over the next 8.5 min. The column was washed at 100 % B for 7 min, then equilibrated to 25 % B for 5 min. Heme and heme biosynthetic intermediates were analyzed by multiple reaction monitoring in positive ionization mode using the following *m/z* transitions, protoporphyrin IX (563.3 to 504.3); heme (616.3 to 557.3); coproporphyrin III (655.3 to 596.3); and coproheme III (708.3 to 649.3). Optimal collision energies, skimmer offset, and tube lens voltages were determined empirically before each experiment. Quantification was performed by comparing analyte AUC values to those of an external calibration line of heme and CP III at 0, 0.3, 1, 3, and 10 μM.

## ACKNOWLEDGMENTS

We thank members of the Skaar Laboratory for critical comments of the manuscript. This work was supported by these following grants from the National Institutes of Health: R01 AI73843 (E.P.S.) and R01AI150701 (E.P.S.) and K99/R00 AI168483 (H.P.). D.L.S. was supported by the Grove City College Swezey Fund and the Jewell, Moore, and MacKenzie Fund. The SEM work was supported by 1S10OD028704-01A1. The Vanderbilt Cell Imaging Shared Resource (CISR) was supported by these following NIH grants: CA68485, DK20593, DK58404, DK59637 and EY08126.

## Author Contributions

H.P. and E.P.S. conceived and designed the experiments. S.M.C. conducted some of the XylE experiments, W.N.B. performed the heme quantification, S.M.C., G.H.H., and D.L.S. created the mutants, E.S.K. conducted the SEM imaging, H.P. carried out all other experiments. H.P. drafted the paper. H.P. and E.P.S. edited the paper. All authors reviewed the paper.

---

**Table S1.** Strains and plasmids used in this study.

| Species | Genotype | Description | Reference |
| --- | --- | --- | --- |
| *B. anthracis*<br>strain<br>Sterne | WT | Wildtype laboratory stock | Lab stock |
|  | 2xRelE (*bas3009::P<sub>hrt</sub>-relE* *bas4599::P<sub>hrt</sub>-relE*) | RelE selection strain (*hrt* promoter fused to relE and inserted into the BAS3009 and BAS4599 pseudogene loci) | (38) |
|  | *fapR::tet* | The *fapR* reading frame replaced with a tetracycline cassette | This study |
|  | *fapR::tet* pOS1.P<sub>lgt</sub>*fapR* | Complementation *fapR* deletion *in trans* | This study |
| | Δ*hemX* | Lacking the *hemX* reading frame | This study |
| | Δ*hemX* pOS1.P<sub>lgt</sub>*hemX* | Complementation Δ*hemX* deletion *in trans* | This study |
| | *fapR::tet* Δ*hemX* | Double mutant of *fapR* and *hemX* | This study |
|  | WT pOS1.P<sub>lgt</sub>*hssS.GFP* | WT harboring a GFP-HssS protein fusion construct. | This study |
|  | *fapR* pOS1.P<sub>lgt</sub>*hssS.GFP* | *fapR::tet* harboring a GFP-HssS protein fusion construct. | This study |
|  | WT pOS1 P<sub>hrt</sub>*XylE* | WT harboring a XylE reporter | This study |
|  | *fapR* pOS1 P<sub>hrt</sub>*XylE* | *fapR::tet* harboring a XylE reporter | This study |
| | *hemX* pOS1.P<sub>hrt</sub>*XylE* | Δ*hemX* harboring a XylE reporter | This study |
| | *fapR hemX* pOS1.P<sub>hrt</sub>*XylE* | *fapR::tet* Δ*hemX* harboring a XylE reporter | This study |
|  | WT pOS1 P<sub>hrt</sub>mcherry | WT harboring an mcherry reporter | This study |
|  | *fapR* pOS1 P<sub>hrt</sub>mcherry | *fapR::tet* harboring an mCherry reporter | This study |
| | *hemX* pOS1.P<sub>hrt</sub>mcherry | Δ*hemX* harboring an mCherry reporter | This study |
| | *fapR hemX* pOS1.P<sub>hrt</sub> mcherry | *fapR::tet* Δ*hemX* harboring an mCherry reporter | This study |
| *E. coli* | DH5α WT | Wildtype laboratory stock for cloning | Lab stock |
|  | K1077 WT | Wildtype laboratory stock for cloning | Lab stock |
| **Plasmid** | **Description** |  | **Reference** |
| pLM4 | Plasmid for gene deletion |  | Lab stock |
| pOS1 | Plasmid for complementation |  | Lab stock |

**Table S2.**
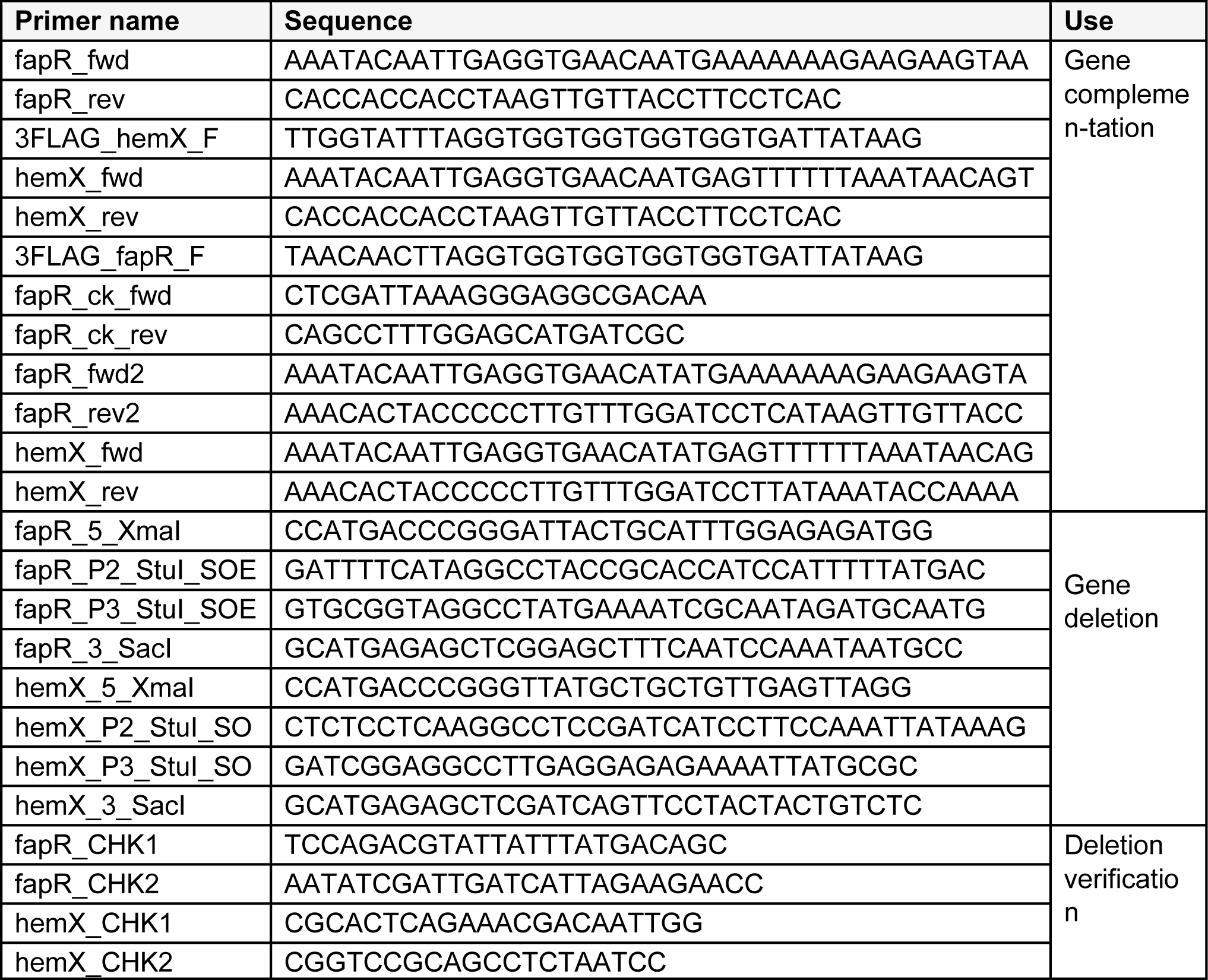
Oligos used in this study.

**Table S3.** Isolated spontaneous resistant suppressors with frameshift mutations in *fapR*.

| Background | Suppressor | Allele | Genotype |
| --- | --- | --- | --- |
| WT<br>*bas3009::P<sub>hrt</sub>-relE*<br>*bas4599::P<sub>hrt</sub>-relE* | S1, S7, S8 | *fapR-1* | Deletion of A at nucleotide 10<br>→ frameshift after amino-acid residue 3 |
|  | S3 | *fapR-2* | Deletion of T at nucleotide 279<br>→ frameshift after amino-acid residue 92 |
|  | S5, S6 | *fapR-3* | Deletion of C at nucleotide 322<br>→ frameshift after amino-acid residue 107 |

**Figure S1.**
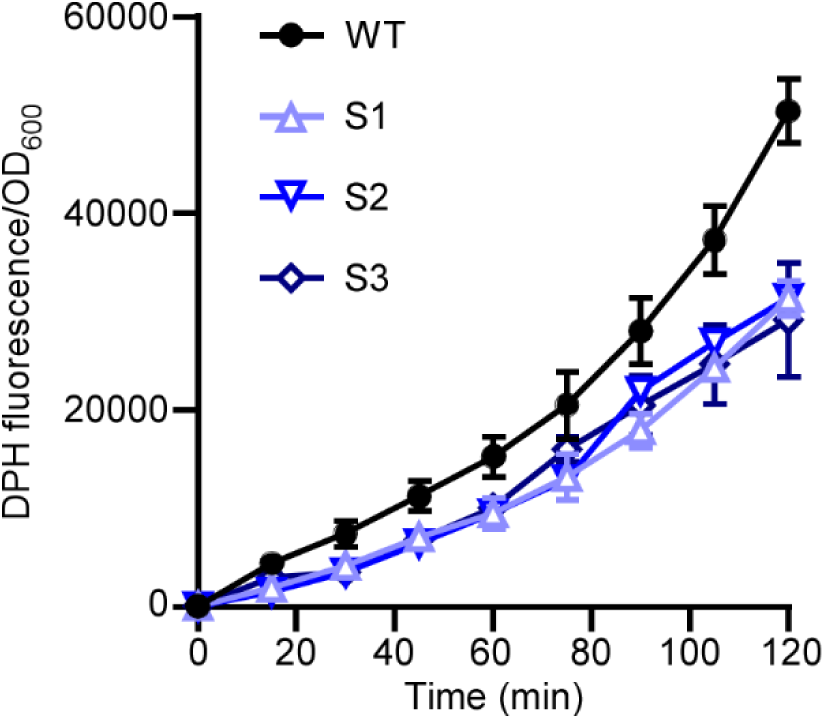
Disruption of *fapR* leads to increased membrane rigidity in *B. anthracis*. Membrane permeability was evaluated over 2 h in *B. anthracis* Sterne WT and the three representative suppressors by monitoring the fluorescence of 1,6-diphenyl-1,3,5-hexatriene (DPH) (excitation, 365 nm; emission, 460 nm). All data represent means ± SD for measurements acquired in three biological replicates.

**Figure S2.**
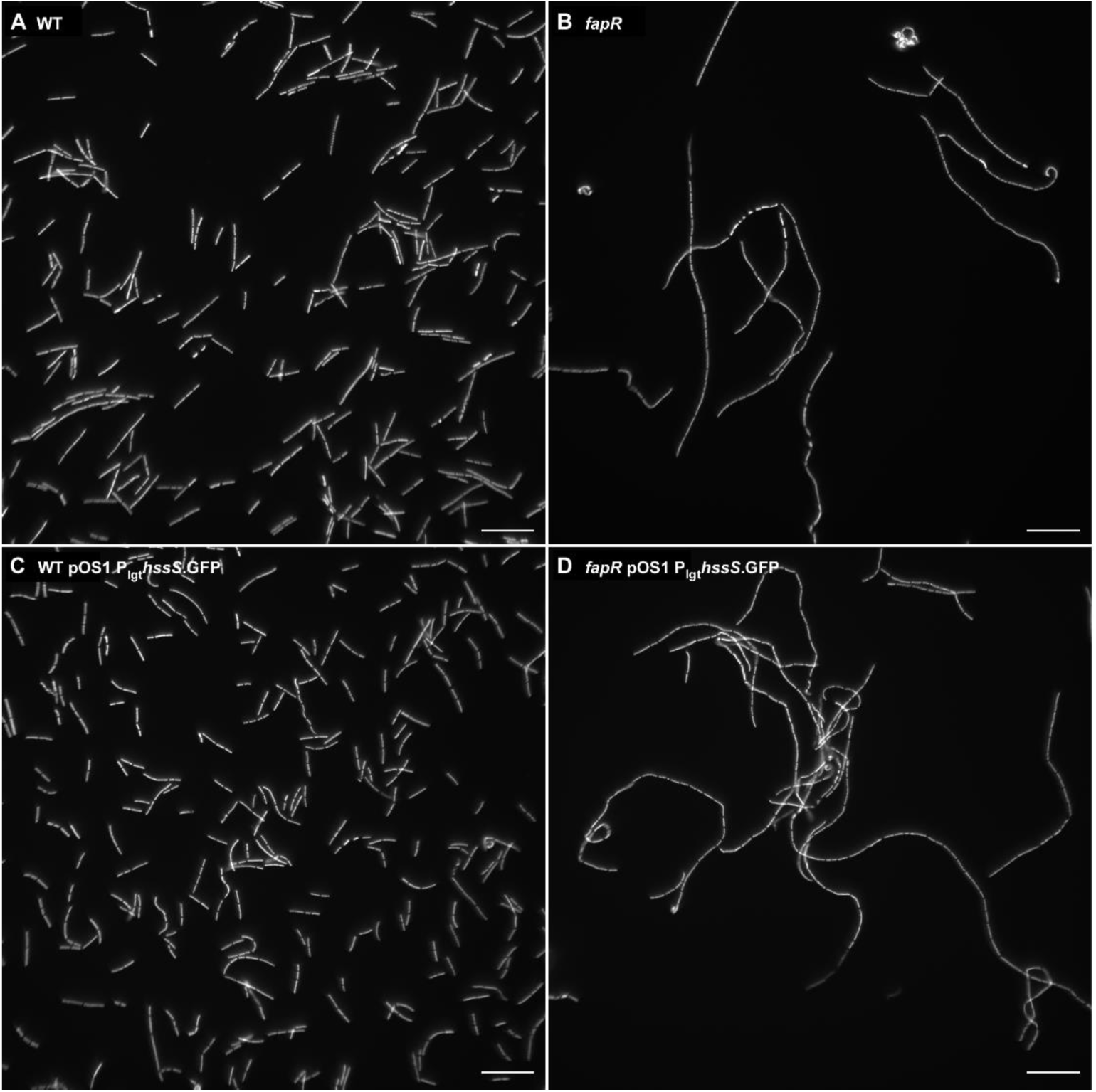
Deletion of *fapR* leads to formation of long curved filaments. Representative fluorescent images of WT, *fapR*, WT pOS1.P_lgt_*hssS*.*GFP*, and *fapR* pOS1.P_lgt_*hssS*.*GFP*. Cells were grown to OD_600_∼1 prior to staining with Hoechst. Scale bar, 20 µm.

**Figure S3.**
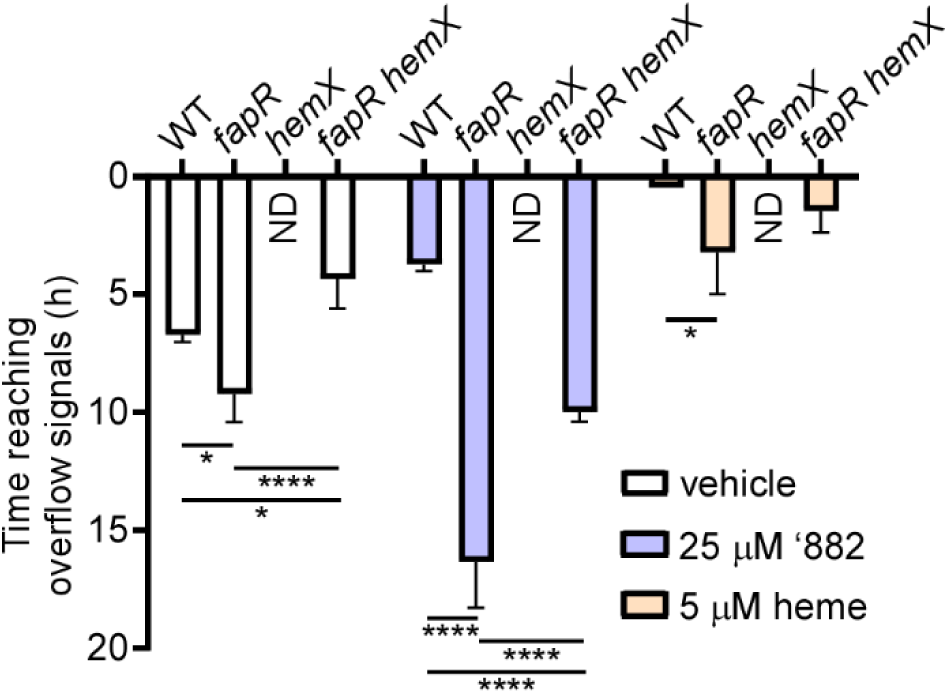
Heme sensing response is highly activated in Δ*hemX* but compromised by deletion of *fapR*. Overnight cultures of *B. anthracis* strains harboring pOS1. P*_hrt_*-*mcherry* were inoculated in LB amended with varied compound as indicated and fluorescence was monitored for 20 h at 37 °C (Ex: 590 nm; Em: 620 nm). The time taken to reach an overflow signal, indicating maximum fluorescence beyond the detection limit, serves as a proxy for P*_hrt_* promoter activation regulated by the HssRS system. The fluorescence signal in the *hemX* mutant reached its maximum instantly so the time required to reach an overflow signal in this mutant is not determined (ND). All data are mean ± SEM (n=9). Statistical analyses were done using two-way ANOVA ∗P < 0.05, ∗∗P < 0.01, ∗∗∗P < 0.001, and ∗∗∗∗P < 0.0001.

